# CODEC enables ‘single duplex’ sequencing

**DOI:** 10.1101/2021.06.11.448110

**Authors:** Jin H. Bae, Ruolin Liu, Erica Nguyen, Justin Rhoades, Timothy Blewett, Kan Xiong, Douglas Shea, Gregory Gydush, Shervin Tabrizi, Zhenyi An, Sahil Patel, G. Mike Makrigiorgos, Todd R. Golub, Viktor A. Adalsteinsson

## Abstract

Detecting mutations as rare as a single molecule is crucial in many fields such as cancer diagnostics and aging research but remains challenging. Third generation sequencers can read a double-stranded DNA molecule (a ‘single duplex’) in whole to identify true mutations on both strands apart from false mutations on either strand but with limited accuracy and throughput. Although next generation sequencing (NGS) can track dissociated strands with Duplex Sequencing, the need to sequence each strand independently severely diminishes its throughput. Here, we developed a hybrid method called Concatenating Original Duplex for Error Correction (CODEC) that combines the massively parallel nature of NGS with the single-molecule capability of third generation sequencing. CODEC physically links both strands to enable NGS to sequence a single duplex with a single read pair. By comparing CODEC and Duplex Sequencing, we showed that CODEC achieved a similar error rate (10^−6^) with 100 times fewer reads and conferred ‘single duplex’ resolution to most major NGS workflows.

## Introduction

Discovering extremely low-level mutations as rare as within a single double-stranded DNA molecule (a ‘single duplex’) is crucial to finding diagnostic[1, 2], predictive[3, 4], and prognostic[5, 6] biomarkers, understanding cancer evolution[7, 8] and somatic mosaicism[9, 10], and studying infectious diseases[11, 12] and aging[13, 14]. Third generation sequencing technologies (e.g., PacBio, Oxford Nanopore Technologies) in principle make it possible to sequence each single DNA duplex in whole to resolve true mutations on both strands apart from false mutations on either strand, but, in practice, lack the required accuracy and throughput[15, 16]. Next generation sequencing (NGS), on the other hand, continues to offer superior read accuracy and throughput[17], but is not configured to sequence single duplexes—at least not without severely compromising its throughput or utility.

NGS affords high throughput by reading short, clonally amplified DNA fragments in massively parallel fluorescence analysis. Yet, its accuracy is limited by the need to dissociate Watson and Crick strands of each DNA duplex. Without a complementary strand for comparison, errors introduced on either strand due to base damage[18], PCR[19], and sequencing[20] can be disguised as real mutations (Fig. 1a). While it is possible to use unique molecular identifiers (UMIs) to separately track both strands of each DNA molecule and compare their sequences to detect true mutations on both strands of each duplex[21, 22], it does not solve the underlying limitation of NGS: duplex dissociation. For example, Duplex Sequencing[23] tags double-stranded UMIs on each original duplex to trace them back after PCR and NGS. By forming a duplex consensus between reads assigned to the Watson and Crick strands of each original duplex, Duplex Sequencing achieves 1,000-fold or higher accuracy (error rate below 10^−6^) and can thus resolve true mutations within single DNA duplexes. However, recovering both strands among up to 10 billion other strands on an NGS flow cell (e.g., Illumina NovaSeq) requires 100-fold excess reads[24], which invariably diminishes the throughput of NGS and severely limits its applicability.

**FIG. 1.**
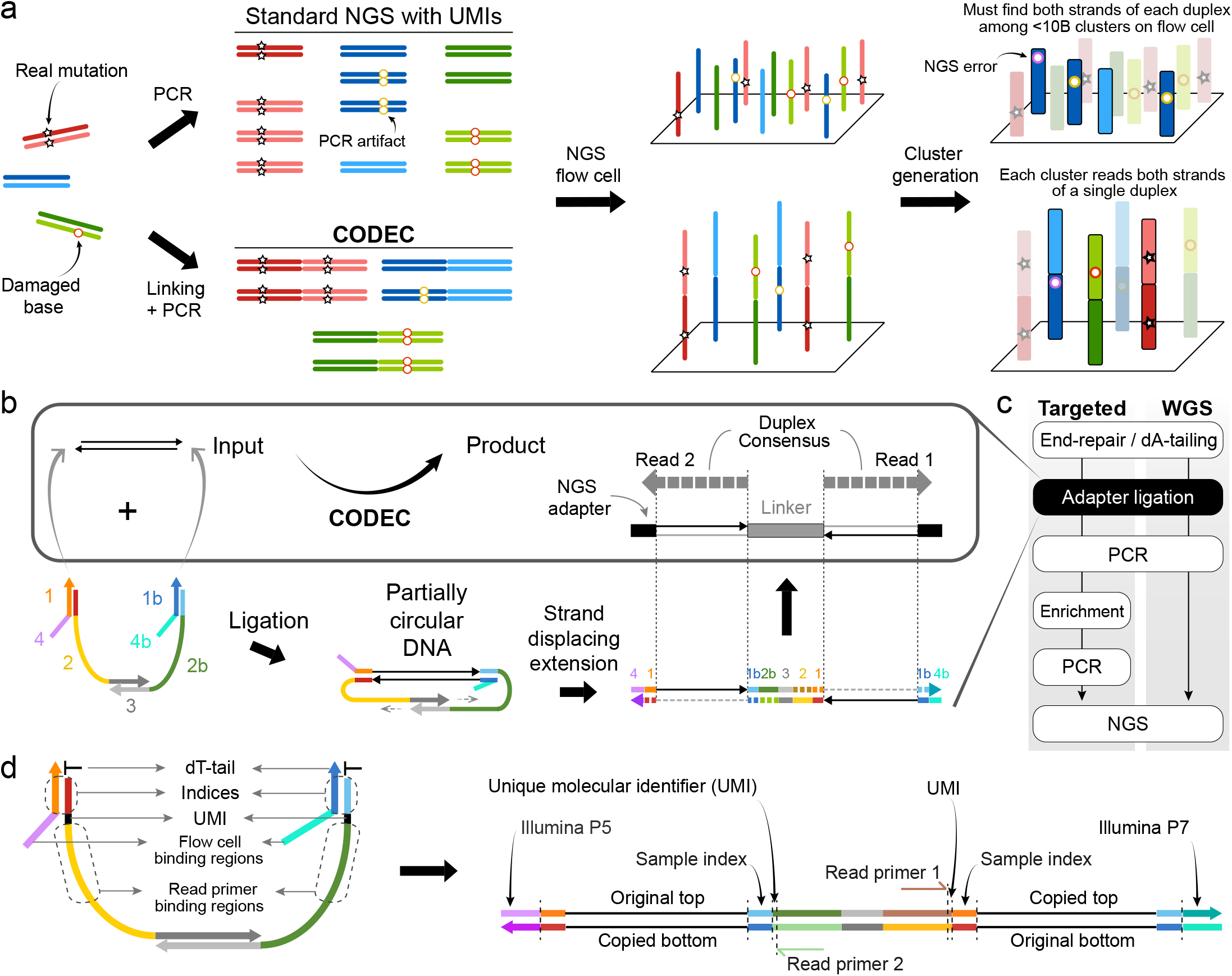
Overview of Concatenating Original Duplex for Error Correction (CODEC). **(a)** Standard NGS workflows involve dissociation of DNA duplex, which loses the intrinsic property of DNA that encodes genetic information twice. Both strands of a duplex can be tracked through unique molecular identifiers (UMIs) to identify false mutations caused by base damage, PCR, and NGS errors, but finding them among <10 billion other strands costs throughput, highlighted by blue clusters. CODEC physically links each duplex before dissociation, ensuring each library molecule retains information of both strands. **(b)** CODEC links the sequence information of an original duplex into a single strand. As a result, each pair of NGS reads becomes self-sufficient for forming a duplex consensus (box). It utilizes the adapter complex instead of a duplex adapter for ligation, followed by strand displacing extension. **(c)** CODEC modifies the ligation step of ligation-based NGS workflows. **(d)** CODEC adapter complex is prepackaged with all of the components needed for Illumina NGS. Unlike standard NGS libraries, CODEC reads outward to sequence a UMI, an index, and an insert together. No indexed primers are required as indices and flow cell binding regions (P5 and P7) are added by the ligation.

To date, a few methods have sought to overcome the high inefficiency of Duplex Sequencing. Duplex Proximity Sequencing (Pro-Seq)[25] uses a polyethylene glycol linker to link 5′-ends of an original Watson strand and a copied Crick strand of a duplex to avoid hairpin formation for whole-genome sequencing (WGS). However, concatenating two strands with the opposite directions blocks DNA amplification which is necessary for most applications. CypherSeq[26] generates a circularized duplex followed by rolling circle amplification, but the lack of asymmetry between the two strands obscures whether both strands were actually sequenced. Some technologies such as o2n-seq[27] and Circle Sequencing[28] are compatible with PCR but only link a single strand of each duplex and thus, lack the ability to create a duplex consensus. BotSeqS[29, 30] uses dilution instead of linking to increase the chance of recovering both strands, but by doing so it only sequences 0.001% of the input DNA. Despite the need for sequencing single duplexes with high accuracy and throughput, there has been no such method with universal applicability. We thus reasoned that linking the information of both strands before dissociation could make NGS capable of reading single DNA duplexes with high accuracy and throughput.

Here, we developed a method that combines the massively parallel nature of NGS and the single-molecule capability of third generation sequencing to sequence both strands of each DNA duplex with single read pairs. In this hybrid approach called Concatenating Original Duplex for Error Correction (CODEC), each molecule becomes self-sufficient for forming a duplex consensus via NGS **(Fig. 1a)**. By using the opposite strand as a template for extension instead of directly linking them, CODEC physically concatenates the sequence information of Watson and Crick strands into a single strand without forming a strong hairpin structure **(Fig. 1b)**. Any differences between concatenated sequences would indicate either non-canonical base pairing created by nucleobase damage or an alteration confined to one strand of the original DNA duplex, or an error introduced during PCR amplification or sequencing. We tested CODEC with different sample types and NGS workflows, and confirmed that it suppressed both single nucleotide variants (SNV) and indel errors as accurately as Duplex Sequencing but with 100fold fewer reads, thereby conferring ‘single duplex’ resolution to NGS.

## Results

### CODEC adapter complex and workflow

The CODEC structure can be built by a streamlined workflow using a commercial ligation-based NGS preparation kit and CODEC adapter complex. First, a typical duplex adapter was replaced with the adapter complex consisting of four oligonucleotides, containing all elements required for NGS. We rationally designed double-stranded segments of the adapter to hold the whole complex based on DNA hybridization thermodynamics **(Supplementary Figure S1a)** and introduced single-stranded segments to mitigate bending stiffness of rigid double helix **(Supplementary Figure S1b)**. After adapter ligation closes both ends of an input molecule, strand displacing extension initiates at remaining 3′-ends to elongate each strand by using the opposite strand as a template. The resulting structure is two original strands concatenated with the CODEC linker in the middle and NGS adapters on both sides. The molecular process depicted in **Fig. 1b** is integrated into the adapter ligation step of commercial NGS library construction kits **(Fig. 1c)**.

To fully utilize the concatenated structure, we also relocated the NGS library components **(Fig. 1d)**. In contrast to the conventional Illumina structure with the NGS read primer binding sites on the outer side, we moved the binding sites to the CODEC linker in the middle and sequenced outward to prevent reading molecules without the linker **(Supplementary Figure S1c)**. Having the binding sites at conventional locations had resulted in poor Quality Scores, which we attributed to template hopping in cluster amplification **(Supplementary Figure S2a)**, whereas moving the binding sites to the linker overcame this issue **(Supplementary Figure S2b)**. Sample indices, which are typically located outer to the read primer binding sites and read separately from the inserts, were moved right next to the inserts. By adding the indices during adapter ligation and reading them with the inserts in a single step, CODEC suppressed index hopping even better than the gold standard of using unique dual indices[31, 32] (0.056% vs. 0.16%). We designed sets of 4 sample indices that collectively have all four bases at every position to ensure high base diversity for proper cluster identification, phasing correction, and chastity filtration **(Supplementary Figure S3)**. Because indexed primers were no longer needed, we were able to include Illumina P5 and P7 segments in the adapter complex and use them as universal primer binding regions.

### Proof-of-concept

We first confirmed that the CODEC workflow could create the intended NGS library structure by converting fragmented human genomic DNA (gDNA) from peripheral blood mononuclear cells into a CODEC-NGS library and sequencing it. Due to the novel structure of CODEC reads, we created a user-friendly analysis pipeline called CODECsuite to process the data (see Methods). We found that more than half of the reads showed the correct structure **(Fig. 2a)**. Meanwhile, the major byproducts appeared to have been created when an input duplex was either ligated to two different adapter complexes (“double ligation”) or no adapter complex (“blank ligation”), or when strand displacing extension occurred between two ligated products (“intermolecular”) **(Supplementary Figure S4)**. Yet, almost 90% of byproducts still retained information on one side of a duplex just like standard NGS, suggesting that the byproducts may still yield useful data.

**FIG. 2.**
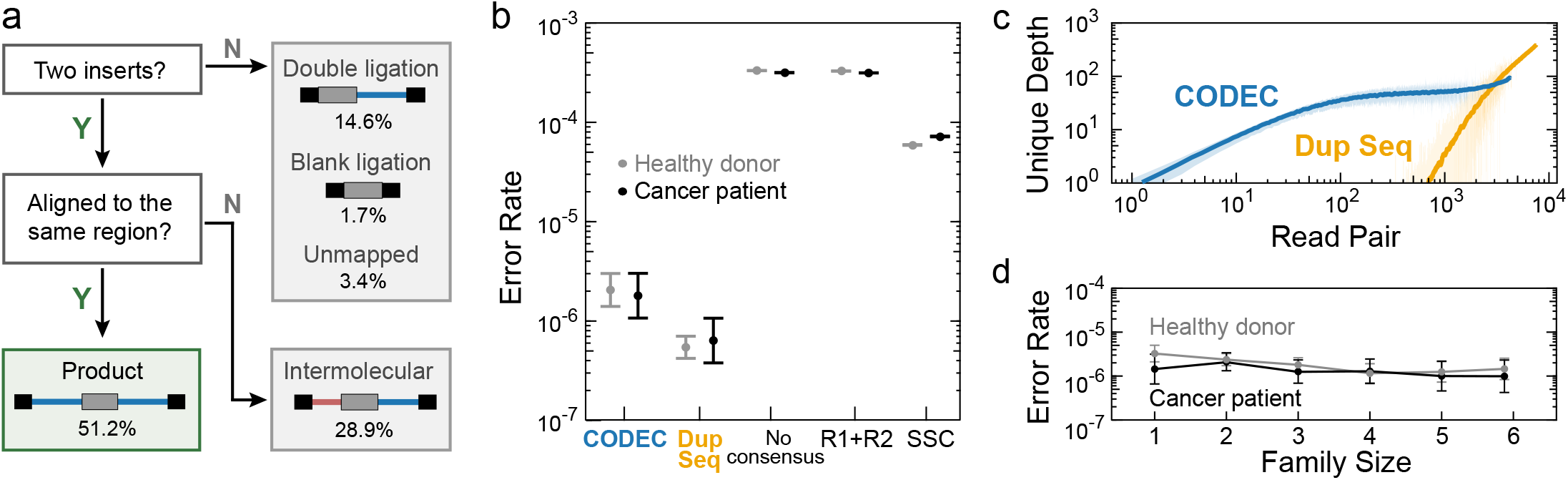
Proof-of-concept. **(a)** Ratios of the correct CODEC product and byproducts which have been named after how they were likely created. **(b)** Error rates of CODEC, Duplex Sequencing, and other consensus methods including typical paired-end read (R1+R2) and single strand consensus (SSC). Target enrichment with a pan-cancer gene panel was performed on cell-free DNA (cfDNA) of two individuals. Error bars indicate 95% binomial confidence intervals. **(c)** Recovery of unique original duplexes per captured region in healthy donor cfDNA against the amount of sequencing. Solid lines show moving averages and shades indicate standard deviations. **(d)** CODEC error rates at each family size, which is the number of raw reads with the same UMI and start-stop positions.

We next explored whether the fragments with the correct CODEC structure could provide comparable error rates to Duplex Sequencing using significantly fewer reads. To assess this, we performed a head-to-head comparison. Because Duplex Sequencing requires high sequencing depth per locus, we ran target enrichment with a pan-cancer panel on NGS libraries prepared with each method, built from 20 ng cell-free DNA (cfDNA) from a cancer patient and a healthy donor. We found that the mean CODEC error rate of two individuals (1.9 x 10^−6^) was similar to that of Duplex Sequencing (5.9 x 10^−7^) **(Fig. 2b)** with no statistically significant difference in sequence contexts of errors except for C:G>T:A in a healthy donor **(Supplementary Figure S5a)**, which we believe could be resolved using an improved end-repair method[30, 33] **(Supplementary Figure S5b)**. Additionally, when error rates were plotted as a function of distance from either end of a fragment, we saw elevated error rates from CODEC and Duplex Sequencing data toward the fragment ends of duplex consensus, consistent with prior reports of error propagation in end-repair[30, 33] **(Supplementary Figure S6)**. This observation reassures that reading a single CODEC fragment is equivalent to reading two Duplex Sequencing fragments from each strand and affirms the need to trim 12 base pairs (bp) from both ends of each original DNA duplex in silico[24].

To further confirm that the error suppression potential of CODEC is uniquely enabled by reading both strands of the original DNA duplex together, as opposed to simply forming a consensus of forward and reverse reads, we then compared error rates of three additional methods from the same NGS data: no consensus, paired-end reads consensus (R1+R2, collapses read 1 and read 2), and single strand consensus (SSC, collapses reads from the same original strand). Interestingly, the error rate gap between the no consensus and R1+R2 was negligible **(Fig. 2b)**, suggesting that many errors are physically present in NGS library molecules, and could have been introduced during library amplification, or when each library molecule undergoes bridge amplification for cluster i generation (Fig. 1a). Although SSC was more accurate than R1+R2 and the no consensus reads, without a consensus of Watson and Crick strands, its error rate was 23-fold higher than that of CODEC. The fact that reading the same strand multiple times does not contribute as much as duplex consensus implies the intrinsic limitation of other sequencing technologies[27, 28].

We next explored the number of reads required to uncover the same number of unique DNA duplexes. When we used UMIs as well as start and stop mapping positions of each molecule to collapse all reads to unique original duplexes, we found that Duplex Sequencing could not start reassembling duplexes until receiving 700 reads **(Fig. 2c)**. In contrast, CODEC started to reassemble 350-fold earlier. The gap between required reads was maximized when recovering a smaller number of duplexes, suggesting that CODEC could be uniquely capable of sequencing broad genomic regions with shallow depth. Notably, even a single paired-end read of CODEC was highly accurate **(Fig. 2d)**, as each CODEC read is self-sufficient to form a duplex consensus. Our results suggest that CODEC confers the accuracy of duplex sequencing from single paired-end reads and thus sequences more DNA duplexes using substantially fewer reads.

### CODEC confers the accuracy of duplex sequencing to WGS and WES

We next sought to determine whether CODEC could enable human whole-exome and whole-genome ‘duplex’ sequencing, which would otherwise be impractical due to high cost. To assess this, we applied CODEC whole-exome sequencing (WES) to gDNA and formalin-fixed paraffin-embedded (FFPE) samples from a cancer patient, whose samples had been tested in our prior publication[24]. We found that CODEC reduced the sequencing error rates of both samples, with 100-fold improvement for gDNA**(Fig. 3a)**. Analyzing the sequence context of the errors revealed that CODEC improved accuracy across all types of SNV **(Fig. 3b)**, suggesting that the capability of CODEC to suppress errors is not limited to specific contexts. Of note, there were more C>T errors in FFPE samples due to deamination artifacts[34], which we believe could be resolved with improved end-repair methods[30, 33].

**FIG. 3.**
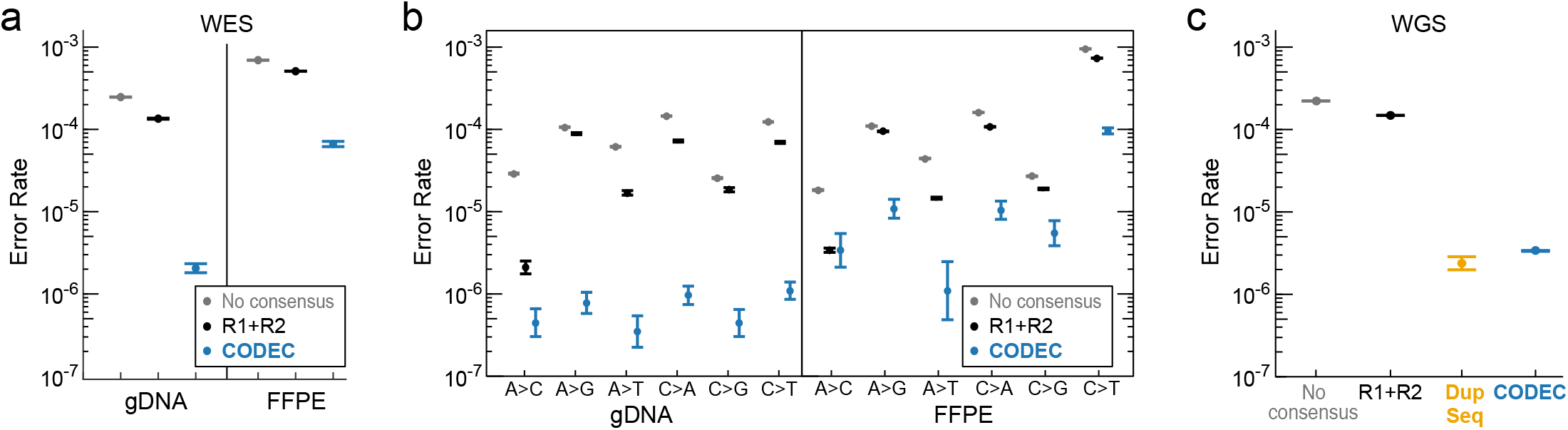
Error rates of whole-exome sequencing (WES) and whole-genome sequencing (WGS). **(a)** Error rates of CODEC on formalin-fixed paraffin-embedded (FFPE) and matching normal samples of a cancer patient. **(b)** Errors in (a) broken down by sequence context. **(c)** Error rates of WGS with Duplex Sequencing and CODEC performed side by side.

Next, we applied CODEC and Duplex Sequencing to WGS of the pilot genome NA12878 of the Genome in a Bottle Consortium (GIAB)[35]. For a fair comparison, we assigned the same amount of sequencing to each method although Duplex Sequencing could not recover many unique duplexes. The error rates of both Duplex Sequencing (2.38 x 10^−6^) and CODEC (3.37 x 10^−6^) were much lower than that of the no consensus reads (2.2 x 10^−4^) or R1+R2 (1.48 x 10^−4^) **(Fig. 3c)**. This result confirms that CODEC is as accurate as Duplex Sequencing under the same conditions. The error rates of each sequence context showed that CODEC has a similar error profile to Duplex Sequencing **(Supplementary Figure S7)**.

Depth of coverage analysis for WGS further demonstrated that CODEC achieved 160-fold greater unique duplex depth than Duplex Sequencing. On the GIAB v3.3.2 hg19 high confidence genomic region (2.6B bases), CODEC had a mean unique duplex depth of 3.96 with 320M raw reads, whereas Duplex Sequencing had only 0.025 mean depth even with 35% more raw read output (431M reads), because most reads did not find their matching strand of the original duplex **(Fig. 4a)**. Thus, we concluded that Duplex Sequencing is not appropriate for WGS and treated Duplex Sequencing WGS data as standard WGS data without generating duplex consensus after this point. In contrast, CODEC covered each base with four unique duplexes on average, confirming the strength of resolving single duplexes.

**FIG. 4.**
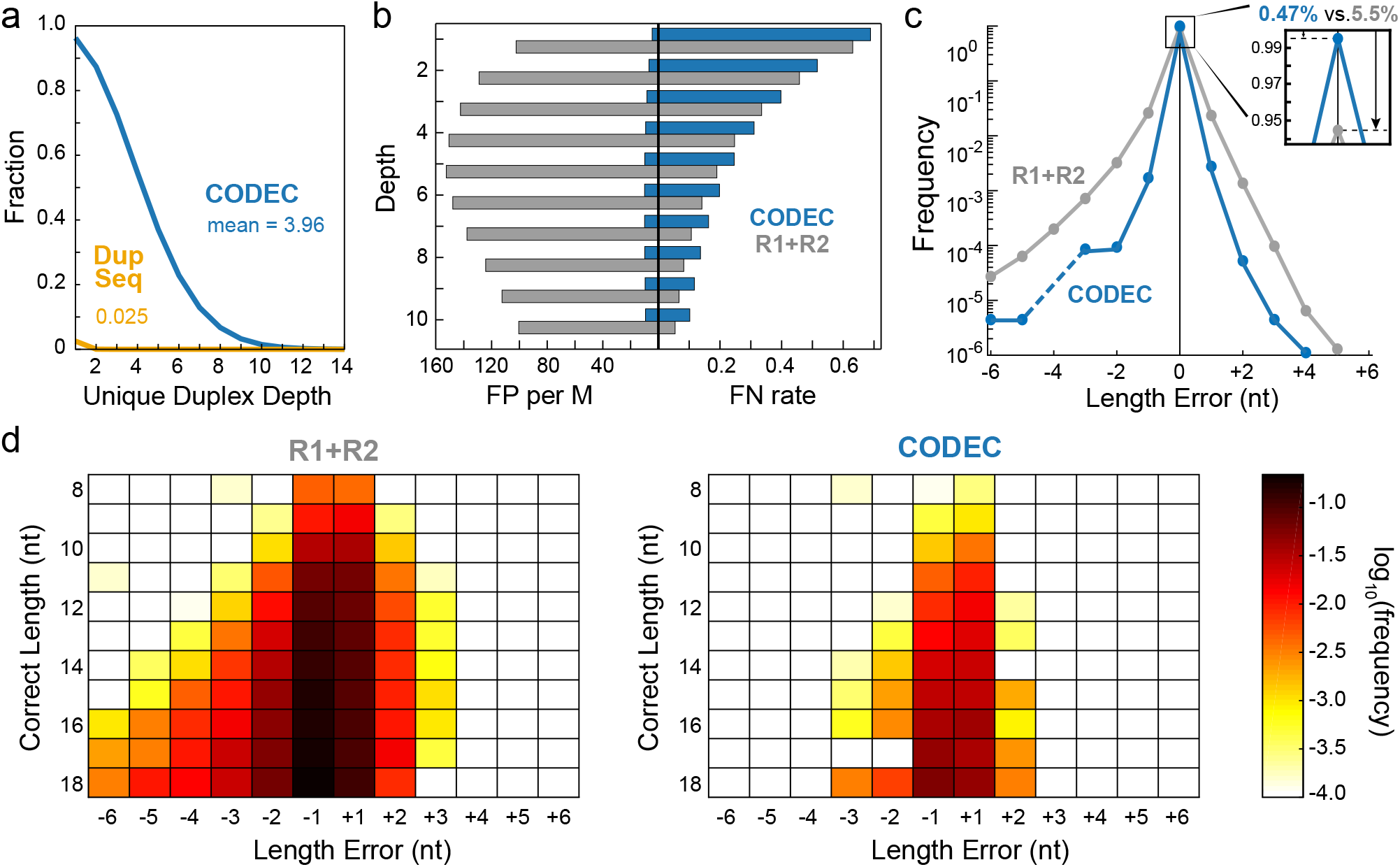
In-depth comparison of WGS results. **(a)** Fractions of each unique duplex depth of CODEC and Duplex Sequencing. **(b)** False positives and false negatives of CODEC and R1+R2 when downsampled to lower depths. **(c)** Summarized indel error frequency at mononucleotide microsatellites. **(d)** Indel error frequency at mononucleotide microsatellites with different lengths from 8 to 18 nucleotides.

### CODEC pushes the frontiers in secondary analysis applications

Achieving the error rate of Duplex Sequencing in WGS/WES gives CODEC the ability to push the limits of many secondary analysis applications. One such application is benchmarking the whole genome small germline variant calling (SNV + indel). To test the potential of CODEC at low coverage as implied in Figure 2c, we compared CODEC data of the aforementioned NA12878 sample against standard NGS (R1+R2) at coverages ranging from 1x to 10x, while acknowledging that state-of-the-art germline calling usually requires 30x depth. GATK4 was used for variant calling and followed by the GIAB best practice for benchmarking small germline variants[35]. CODEC showed 90% fewer false positives (FP) than standard WGS with R1+R2 at a cost of 5% higher false negatives (FN) across all downsampled depths **(Fig. 4b, Supplementary Table S1)**. By downsampling NGS data, we also observed how FP and FN are affected by the depth. The lower level of FP in CODEC was the expected result, considering its lower error rate. Its FN levels were slightly higher than that of standard WGS, probably because the lower library conversion efficiency resulted in higher duplication rate, but the difference between FN rates of CODEC and standard WGS became smaller as the coverage decreased. Meanwhile, the advantage of having low FP became more significant at the lower coverage, implying that applications with shallow depth could benefit more from using CODEC.

Considering CODEC’s performance for indel detection at low coverage, we thought that CODEC could improve the sequencing accuracy of microsatellites (MS), which are well-known mutation hot spots. Indeed, when the reference sequences of the MS in NA12878 were compared between CODEC and standard NGS results, CODEC showed lower frequencies of both insertion and deletion errors than standard WGS at mononucleotide MS from 8 to 18 nucleotides **(Fig. 4c)**. The ratio of CODEC reads with incorrect MS lengths was 0.47%, which was 12 times lower than that of standard WGS. Such lower frequencies were consistently observed across mononucleotide MS of varied lengths **(Fig. 4d)** These findings imply that CODEC could be used to read the repeat numbers/copy numbers of MS sites for detecting microsatellite instability (MSI). MSI has been shown to be a predictive marker of response to cancer immunotherapy but remains challenging to detect at low frequency such as from liquid biopsy samples[36]. Tracing mutations in MS is also useful for tracing cell lineages and evolution[37]. The improvements in the secondary applications we have shown highlight what CODEC could enable by sequencing a single duplex within each NGS cluster.

## Discussion

By physically linking both strands of each DNA duplex, CODEC enables each NGS cluster to have single duplex resolution like third generation sequencers. Unlike Duplex Sequencing which requires dissociating duplexes and recovering them back to form a duplex consensus, CODEC distinguishes real mutations from errors with similarly high accuracy but with 100-fold fewer reads. We first showed the proof-of-concept of our approach using cfDNA enriched by a pan-cancer panel, followed by testing its consistency across other major NGS workflows (e.g., WES and WGS) and sample types (e.g., FFPE and germline DNA). To present more uses of CODEC, we also showed that it suppressed FP especially at shallow sequencing depth and reduced indel errors at MS sites.

In a head-to-head comparison, we showed that CODEC is as accurate as Duplex Sequencing but with a much lower sequencing requirement, which has been a major limitation of Duplex Sequencing. Because an error rate is affected by multiple factors other than a sequencing technology itself, any direct comparison requires everything else to be the same. We used the same experimental and computational protocols whenever applicable, including input samples and mass, reagents, target regions, definition of an error, and analysis pipelines for precise comparison.

Because CODEC redefines standard NGS with a novel molecular structure, there may still be room for improvement in its use with target selection protocols including hybrid capture, multiplexed amplicon, and mutation enrichment sequencing[38]. We are also working to improve CODEC’s conversion efficiency. The CODEC adapter complex is attached through two consecutive ligations: a bimolecular ligation followed by a unimolecular ligation. Unlike typical bimolecular adapter ligation where increasing adapter concentration also increases conversion efficiency, unimolecular ligation could be less favorable when the adapter concentration is too high. Consequently, the current version of CODEC adapter complex needs balancing between two ligations. We are currently developing another version of CODEC that circumvents two consecutive ligations.

Although conventional end-repair/dA-tailing of a commercial kit was used throughout this work, the accuracy can be further improved if a new end-repair method is adopted before CODEC. Recent studies[30, 33] have reported that base damage on overhangs and single-stranded breaks of original DNA duplexes can lead errors on one strand to be copied to both strands. It was also indirectly observed in this work that error rates were generally higher toward the ends of DNA fragments (Supplementary Figure S6). While such errors appear on duplex consensus and result in false mutations, new end-repair methods prevent the error propagation, and we believe that even higher accuracy will be attainable when CODEC is combined with new end-repair methods[30, 33].

Reading a single CODEC fragment is equivalent to reading both strands of an original duplex, which eliminates the need to read the same locus multiple times. The low error rate of CODEC at 1x read depth opens possibilities for various applications across fields from diagnostics to bioinformatics. One example is discovering rare somatic mutations with a limited number of reads, which has a higher chance of finding a true mutation when the error rate gets lower[39]. Another example is shotgun metagenomic sequencing for microbiome analysis, where suppressing false SNVs with CODEC would prevent incorrect taxonomic classifications and inaccurate evaluation of microbial diversity[40]. In de novo assembly, lower error rates contribute to more contiguous assembly in de Bruijn graph paradigm and faster process in overlap-layout-consensus paradigm[41].

In summary, CODEC transforms standard NGS instruments into massively parallel ‘single duplex’ sequencers by concatenating both strands of each original DNA duplex. This strategy enables SNV and indel detection as accurate as Duplex Sequencing, even in cases where Duplex Sequencing is not possible due to low throughput. We thus believe that CODEC could be broadly enabling for many important biomedical applications such as detecting early-stage cancer or minimal residual disease from liquid biopsies, clinically actionable mutations from liquid or tumor biopsies, clonal hematopoiesis of indeterminate potential (CHIP) from blood samples, somatic mosaicism in normal tissue samples, and beyond.

## Methods

### DNA samples and oligonucleotides

Cell-free DNA of patient 315 from cohort 05-246 and both FFPE and gDNA of patient 95 from cohort 05-055 were from another study[24]. NA12878 was purchased from Coriell. All samples were stored in low TE buffer (10 mM Tris-HCl, 0.1 mM EDTA, pH 8) and were fragmented by Covaris ultrasonicator to have a mean size of 150 bp except cfDNA. All oligonucleotides for CODEC were synthesized by Integrated DNA Technologies (IDT) and went through PAGE purification (See **Supplementary Table S2** for their sequences). The adapter for Duplex Sequencing was custom-ordered for the Broad Institute by IDT.

### CODEC

The CODEC adapter complex was prepared by diluting four 100 *μ*M oligonucleotides to 5 *μ*M with low TE buffer and 100 mM NaCl, followed by heating at 85 °C for 3 minutes, cooling with −1 °C/min to 20 °C, and incubating at room temperature for 12 hours. Mastercycler X50 (Eppendorf) and MAXYMum Recovery PCR tubes (Axygen) were used for the annealing. The annealed adapter complex was kept at −20 °C for future use. We used NEBNext Ultra II DNA Library Prep Kit for Illumina (New England Biolabs, NEB) and followed the manufacturer’s manual with several exceptions:

1. ligation time was increased to 1 hour, 5 *μ*M adapter complex was diluted with adapter dilution buffer (10 mM Tris-HCl, 1 mM EDTA, 10 mM NaCl, pH 8) to 500 nM before use and replaced NEB adapter,
2. 3 *μ*L of 5′-deadenylase (NEB) were added to ligation reaction,
3. strand displacing extension (sample 40 *μ*L, 10x buffer 10 *μ*L, 0.2 mM dNTP, polymerase 1 *μ*L, H_2_O up to 100 *μ*L) was performed with phi29 DNA polymerase (New England Biolabs) at 30 °C for 20 minutes, followed by standard AMPure XP (Beckman Coulter) clean up with 0.75x volume ratio,
4. KAPA HiFi HotStart ReadyMix and xGen Library Amplification Primer Mix (IDT) were used for PCR by following the manufacturer’s manuals with 2 minutes of extension,
5. and AMPure XP clean up with 0.75x volume ratio was performed twice after the PCR.

Libraries for standard NGS and Duplex Sequencing were prepared as described elsewhere[24]. All Library preparations were performed on twin.tec PCR Plates LoBind 250 *μ*L (Eppendorf). Library quantitation was performed with Qubit dsDNA HS kit (Invitrogen) paired with Bioanalyzer DNA High Sensitivity chips (Agilent).

### Enrichment

Both pan-cancer and WES enrichment was performed with xGen Hybridization and Wash kits and xGen Blocking Oligos (IDT), following the manufacturer’s manual. For capture probes, xGen Pan-cancer Panel (IDT, 800 kb) and custom WES panel for the Broad Institute by Twist Bioscience were used.

### Sequencing

Standard NGS and Duplex Sequencing were performed with Illumina HiSeq 2500 Rapid Run (300 cycles) for a pan-cancer panel and WGS. CODEC was performed with Illumina HiSeq 2500 Rapid Run (500 cycles) for a pan-cancer panel and WGS, and NovaSeq SP (500 cycles) for WGS and WES. The extra cycles were used to confirm the CODEC structure

### CODEC data processing

Due to the unique CODEC read structure, we developed CODECsuite (available at https://github.com/broadinstitute/CODECsuite) to process CODEC data **(Supplementary Note)**. CODECsuite is written in C++14 and python3.7 and we use snakemake6.0.3[42] as the workflow management system. CODECsuite consists of 4 major steps: demultiplexing, adapter trimming, consensus calling and computing accuracy. The first 3 steps are specific to CODEC data. The workflow also involves other standard tools such as BWA[43], Fgbio and GATK[44]. Illumina bcl2fastq was used to generate fastq files (with -R -o, no -sample-sheet because CODECsuite will demultiplex), but is not included in the suite. To speed up the data processing, we recommend splitting the fastq files in batches and processing them in parallel. In this study, using 40 batches, the preprocessing (demultiplexing and adapter trimming) of 800M NovaSeq reads took just a few hours in a HPC environment where each batch was executed using a single CPU and 8G RAM. After demultiplexing and adapter removal, we mapped the raw reads using BWA(0.7.17-r1188) against human reference hg19. Fgbio (https://github.com/fulcrumgenomics/fgbio) was then used to collapse the PCR duplicates and to form essentially single-strand consensus (SSC) reads. These SSC reads were then mapped to the reference genome using BWA again. Next, the duplex consensus reads between R1 and R2 were generated from the SSC alignments. We filtered a consensus base if any of the bases from R1 or R2 has base quality less than 30. The duplex consensus reads were aligned to the reference genome using BWA and the subsequent alignments were indel realigned using GATK3 (https://hub.docker.com/r/broadinstitute/gatk3).

### Duplex Sequencing data processing

Duplex Sequencing data processing used in this study has been described elsewhere[24, 38]. Briefly, Fgbio was used to generate duplex consensus and to filter the consensus reads. The entire workflow and more details are available at the CODECsuite github. Read families with at least 2 copies of each strand were required for generating duplex consensus except for Duplex Sequencing WGS, which relaxed the requirement to 1 copy of each strand to get the best possible duplex recovery.

### Duplex recovery and downsample to certain family sizes

Two custom python scripts were used to generate Figure 2c and 2d, respectively. For duplex recovery, we subsampled the pre-consensus family-assigned reads (after Fgbio GroupReadsByUmi) per target at log spaced fractions starting from 10^−4^ (np.logspace(−4, 0, 30)) and calculated the number of duplex formed at each downsample fraction. In this study, this allowed us to understand situations when only limited sequencing was given (e.g., < 100 read pairs). To understand the impact of family size on error rate, we wrote another python script for downsampling. In our sample, the number of duplex consensus having the exact family sizes (number of pre-collapsed raw reads) were limited and thus gave less confident results. Thus, we took advantage of families with strictly larger family sizes and downsample them to the target family size. We also sought to maintain an equal or close ratio between the number of reads from each strand.

### Error rates in capture sequencing

Throughout the article, we defined the error rate as substitution error rate at the base level after mapping to the reference genome (hg19). We used the substitution error rate for calculating the general error rates because Illumina sequencers usually generate 100-fold less indel errors[45] and this definition is compliant with what other studies have reported[30]. For panel sequencing with match normal, we used Miredas to calculate the error rate in concordance with our previous work[24]. The duplex BAMs from both cfDNA and matched normal samples were generated in the same way and were applied to the same set of filters: 1. no secondary and supplementary alignments; 2. Mapq ≥60; 3. Levenshtein distance (L-distance) between the reads excluding soft clipping and reference genome ≤5 and number of non N-base L-distance ≤2; 4. Excluding bases within 12 bp distance from both fragment ends. In order not to confuse errors with real mutations, we pre-computed the germline SNVs and using GATK4 (HaplotypeCaller[46]) from the Duplex Sequencing normal samples as they have higher on-target ratio and hence coverage (89% vs 40% of CODEC). For the patient sample, we found three somatic SNVs (median VAF=0.26, range 0.24 - 0.28) in the captured regions **(Supplementary Table S3)** using MuTect[39]. Those somatic mutations (patient sample only) and germline mutations were masked when calculating the error rates. The error rates were only reported for cfDNA samples and the match normal were used for filtering possible germline (failed to call or did not pass quality filter by HaplotypeCaller) and CHIP. Thereby we also masked any SNV positions where there were at least 1 duplex read support in match normal samples as CHIP can occur at very low mutation frequency. Finally, the specificity checks[24] were performed on cfDNA samples to remove substitutions that may rise from alignment errors.

### Error rate in whole genome sequencing

The WGS error rate was computed similarly to capture data, except for a few differences. 1, We used ‘codec accuracy’, a C++ program, as a replacement for Miredas due to its speed improvement. 2, We used v3.3.2 GIAB NA12878 high confidence VCF and BED[35] file as germline masks and evaluation regions. 3, there was no match normal. 4, we forwent specificity checks as it is also very slow for large genomes.

### Germline SNV and small indel calling in downsampled WGS

We merged the HiSeq 2500 Rapid Run and NovaSeq SP CODEC data to evaluate germline variant calling. The merged CODEC and standard WGS NA12878 samples were downsampled to 1 to 10x (step size 1x) median coverage in the high confidence regions using GATK DownsampleSam. Next, we ran GATK4.1.4.1 best practices pipeline via Cromwell and Terra workflow (available at web resources) and computed on the Google Cloud Platform. We used RTG vcfeval to calculate False Positives (FP) and False Negatives (FN) for SNVs and indels (< 50 bp) without penalizing genotyping error (if heterozygous variants are called as homozygous and vice versa) using v3.3.2 high confidence VCF and BED file as input. We then calculated FP per million bases by normalizing against the high confidence region size and FN ratio by dividing FN by the total number of true variants.

### Microsatellite instability detection

The full-coverage CODEC consensus BAM and full-coverage standard NGS R1R2 consensus BAM on NA12878 were compared against each other to demonstrate CODEC ability to correct PCR stutter errors and thus to reduce background noise for MSI detection. MSIsensor-pro[47] was used to scan the hg19 for homopolymers of size 8 - 18 nt. Since MSIsensor-pro does not have mapping quality or secondary alignments filters, we pre-filtered the BAM using SAMtools[48] by requiring mapq ≥60 and no secondary or supplementary alignments. And then it was used again to count the number of reads that support different lengths of homopolymer at those pre-selected sites. We removed any homopolymer sites that overlap or are in close proximity (+/−5 bp) with any germline variants. After that, the reference lengths of the homopolymer sites were considered as true lengths. And observed length distributions from reads were compared against truth. The results were generated from chromosome 1 only.

## Code availability

CODECsuite and examples and tutorials including how to regenerate the figures in the manuscript are available at the github site https://github.com/broadinstitute/CODECsuite. The end-to-end workflow is available at https://github.com/broadinstitute/CODECsuite/tree/master/snakemake.

## Data availability

CODEC data and Duplex Sequencing data will be available on dbGAP.

## Acknowledgements

The authors acknowledge the Gerstner Family Foundation for its generous support. This study was also supported in part by SPARC award from the Broad Institute.

## Author contributions

Conception and design: V.A.A., J.H.B.

Method development: V.A.A., J.H.B., R.L.

Data acquisition: J.H.B., E.N.

Data analysis: R.L.

Data interpretation: V.A.A., Z.A., J.H.B., T.B., G.G., R.L., E.N., S.P., J.R. D.S., S.T., K.X.

Writing and review of the manuscript: V.A.A., J.H.B., T.R.G., R.L., G.M.M.

