## Supplementary Figures and Notes for "CODEC enables ‘single duplex’ sequencing"

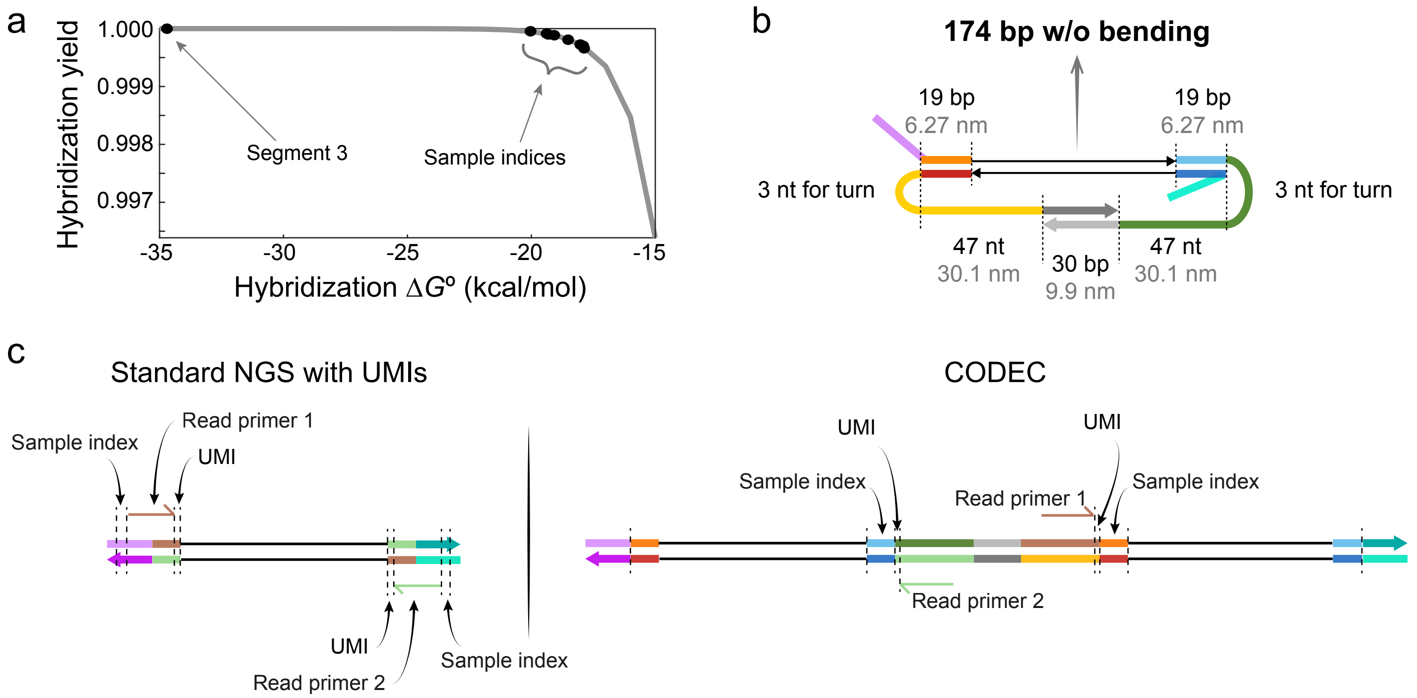


FIG. S1. **Theory behind CODEC adapter complex design.** **(a)** Double-stranded regions of the adapter are predicted to stay stable with oligonucleotide concentrations of 500 nM at 20 °C and $[\mathrm{Na}^+] = 10$ mM. **(b)** The length of single-stranded linkers (yellow and green) was determined to mitigate bending stiffness of a target duplex. The length of dsDNA is shorter along its axis compared to when it is single-stranded. Duplexes with up to 174 bp can be accommodated without bending at all. **(c)** Read primer binding sites of standard NGS and CODEC.

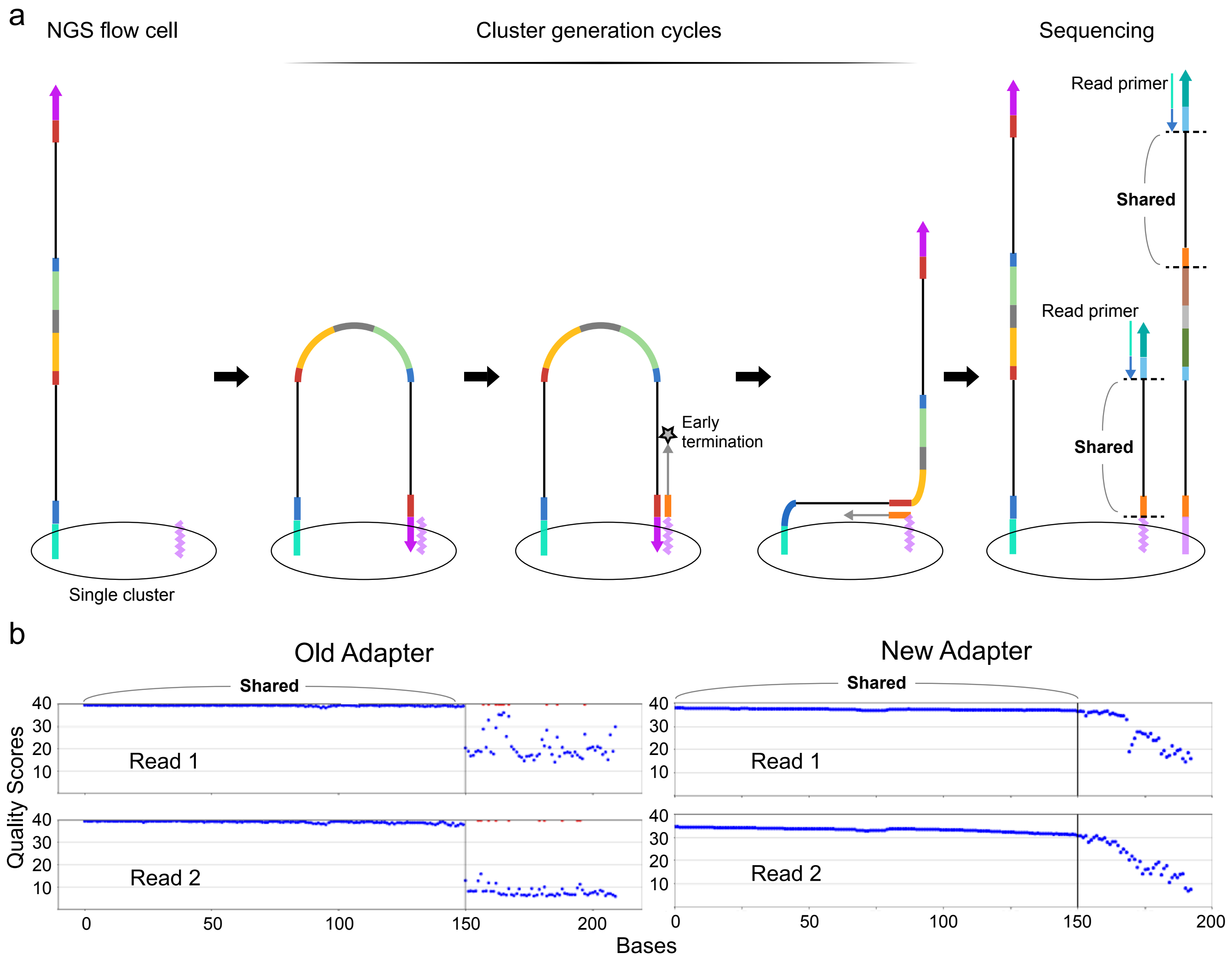


FIG. S2. **(a)** During cluster generation cycles on an NGS flow cell, early termination in the middle of the insert region could create byproducts which turn into shorter fragments with only one insert and the read primer binding regions. These subclonal fragments have the same sequence as the correct fragments until the shared region ends. After sequencing cycles pass the shared region, the short fragments cause mixed fluorescence, and consequently, low Quality Scores. **(b)** Mean Quality Scores of each sequencing cycle by taking the last 150 bp in the shared region and the first 50 bp after the shared region from randomly selected 100 read pairs. Before redesigning the adapter structure, Quality Scores suddenly dropped after the shared region, making it difficult to confirm whether a read has the CODEC structure or not. This issue was solved by moving the read primer binding regions to the linker in order to 'silence' all byproducts without the linker.

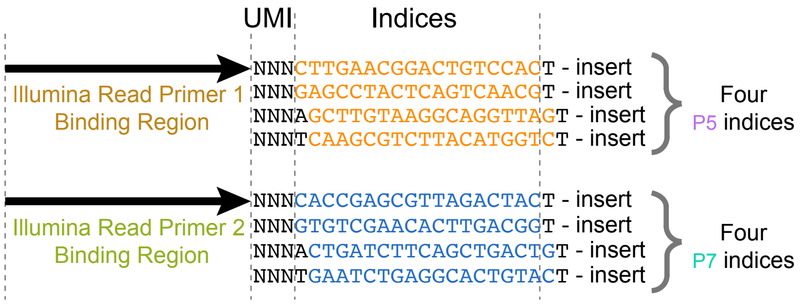


FIG. S3. UMIs and each set of 4 indices are designed to collectively include all four bases at each position while keeping similar hybridization $\Delta G^\circ$ (Fig. S1a) for high-quality image analysis of Illumina sequencers. For example, Illumina software uses up to first 25 bp for various purposes such as cluster identification, phasing correction, and chastity filter.

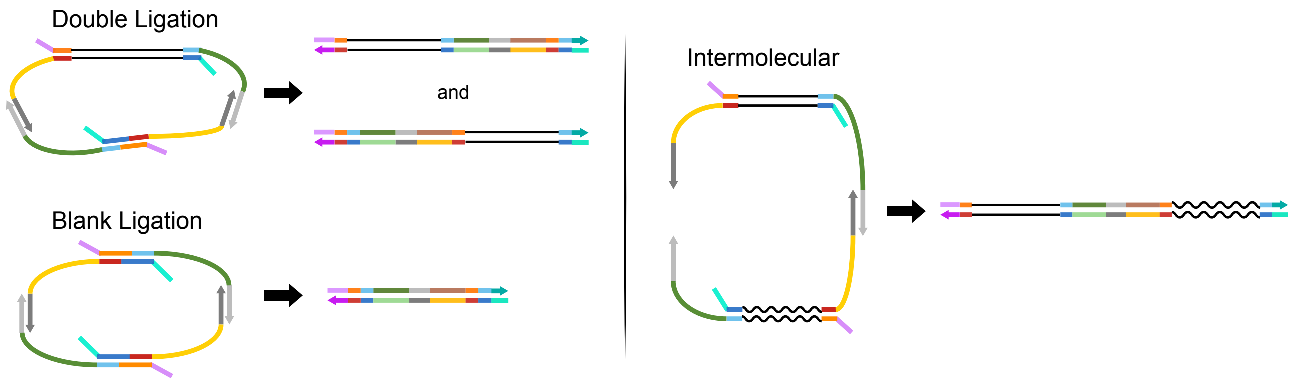


FIG. S4. Expected mechanisms of byproduct formation. "Double ligation" can occur when two adapter complexes are ligated to each end of an insert and go through T/T mismatched ligation with each other, as opposed to A/T ligation. "Blank ligation" can occur when two adapter complexes go through T/T mismatched ligation on both ends with each other with no insert. "Intermolecular" can occur when a strand displacing extension uses another ligation product as a template instead of the opposite strand.

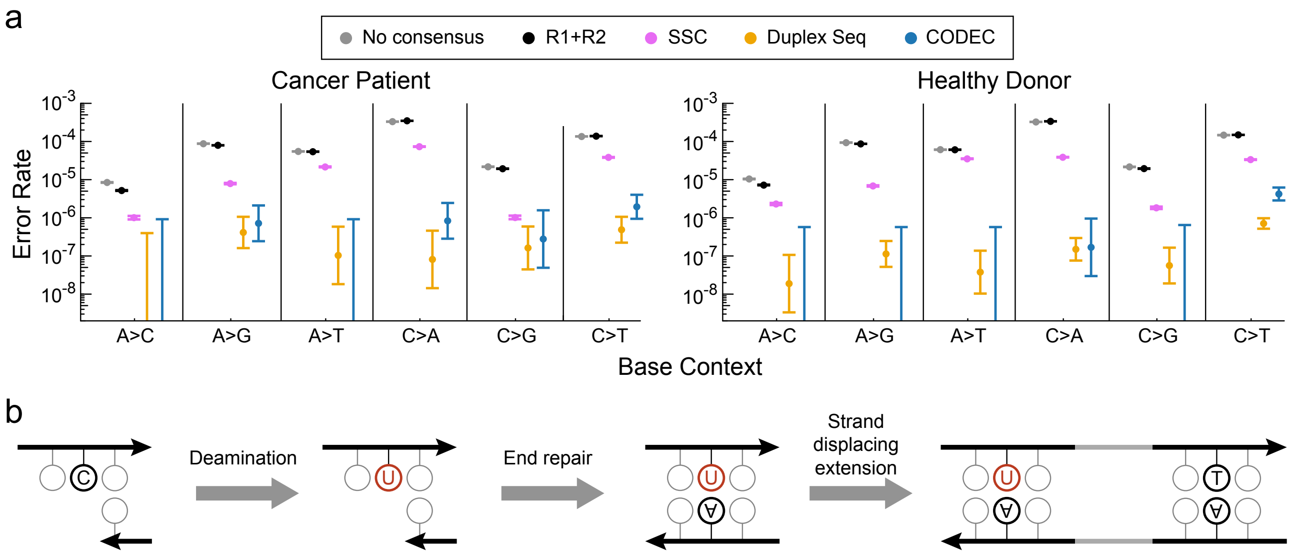


FIG. S5. **(a)** Error rate per base context of targeted sequencing. C>T error rate of CODEC from a healthy donor was higher than that of Duplex Sequencing. **(b)** Possible explanation is that deaminated cytosines, which are uracils, on overhangs of input samples went through end-repair and strand displacing extension. Phi29 DNA polymerase used for the extension can recognize uracils unlike HiFi polymerases and may have created a strand that can be amplified in a subsequent PCR (Crick strand). In case a sample has a high level of deamination, we added USER enzyme step to CODEC workflow in order to suppress false positives from uracils.

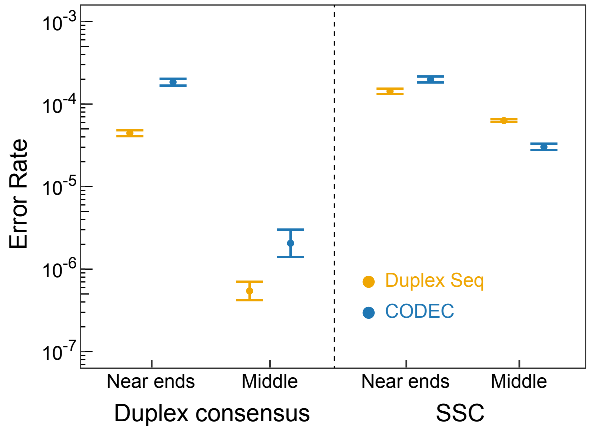


FIG. S6. In duplex consensus data, higher mean error rates of 12 bp from both fragment ends than those of the middle regions imply base damage at 5′-overhangs before end-repair, which was previously observed in other studies using Duplex Sequencing. This is because end-repair fills in 5′-overhangs and copies damaged bases on one strand to both strands and creates false duplex consensuses. In contrast, SSC corrects base damage at neither overhangs nor duplex regions, and thus, shows less error rate differences between the last 12 bp and the middle regions.

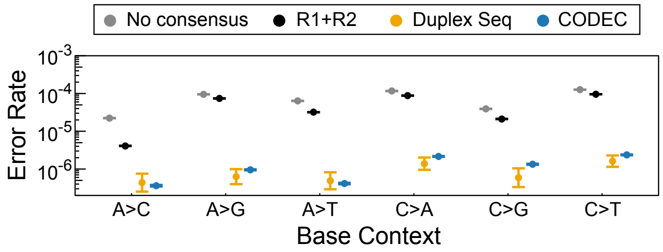


FIG. S7. Error rate per base context of WGS.

### Supplementary Notes: CODECsuite

#### Demultiplexing

CODEC sequencing reads start with Unique Molecular Identifier (UMI) sequences: NNN or NNNA or NNNT (NNN is a random 3-mer), and follow by an 18 bp sample barcode and then a T base (Supplementary Figure S3). To demultiplex, CODECSuite extracts the barcode (4th - 21st bases from the 5′-end) and uses smith-waterman (SW) algorithm[1] for sample indices (SID) assignments. If the extracted barcode is within x edit distance (default 3) away from one and only one sample index, it is declared as a match. Then, a read pair is successfully demultiplexed if and only if the two extracted barcodes (one from each end of the read pair) both match the expected SID (P5 and P7). Only successfully demultiplexed reads are used for subsequent steps and the expected SID are stored in the read names for the subsequent adapter trimming step. Besides, when the two barcodes from a read pair match a chimeric sample index combination, CODECsuite also checks index hopping by aligning the two inserts and flags them as hopping reads if they overlap. Otherwise, the mixed indices are most likely a result of intermolecular byproduct.

#### Adapter trimming and byproducts cleaning

The demultiplexing step adds SID to the read name but does not alter the read sequencing. The adapter trimming step removes the adapter sequences from the read and output as uBAM (unmapped BAM format). The first 3 bases of R1 and R2 are cut and hyphenated and added to the ‘RX’ tag in the bam record. Each correct CODEC read contains a 5′adapter and a possible 3′ adapter (in sequencing orientation). The R1’s SID is used as the template to trim the R1’s 5′ adapter and the reverse complement of R2’s SID is used to trim R1’s 3′ adapter, and vice versa for trimming R2. Again, SW algorithm is used to find a match. We group the reads based on if the 5′ adapter is found on both R1 and R2. In other words, only read pairs with 5′ adapters found in both are considered as potential correct reads. However, a few byproducts can also satisfy this criterion. Therefore, it is important to check the 3′ adapter, if it exists. If a 3′ adapter is found and the insert part is too small (e.g., $<$ 15 bp), we discard the read. If both R1 and R2 are discarded, this template is considered as a blank ligation. If only one of the read ends is discarded, it is classified as a double ligation. The summary of byproducts formation and quantification is made by a custom python script also available at the CODECsuite github site.

#### ReadPair/Duplex consensus

CODECsuite can generate de novo or reference-based consensus. The reference-based consensus has better accuracy and is used throughout this study. A consensus base is formed if two aligned bases (or gaps in terms of insertion or deletion) agree and N otherwise. CODECsuite keeps the pair-end reads but replaces the read sequence with consensus sequence for both R1 and R2. The sequence quality and other auxiliary tags such as UMI are kept intact. The consensus is generated at uBAM format.

#### Alignment accuracy

CODECsuite provides a handy and fast tool for evaluating base level accuracy after alignment. It evaluates bases within bed file regions (such as GIAB high confidence regions) and masks against variants in the VCF and/or MAF file, usually for germline variants and somatic variants respectively. It filters at read level (e.g., mapq or edit distances) and base level (by base quality). It also provides abilities to trim from both fragment ends, and evaluates only the overlapping part of the paired reads. It computes accuracy on fragments, cycle and sample levels. For all non-reference bases, it can output details such as base substitutions, quality score, positions on read and reference so that a post processing script can generate error rate by monomer context.

[1] Smith, T. F. & Waterman, M. S. Identification of common molecular subsequences. J. Mol. Biol. 147, 195–197 (1981).
