## Supplementary Tables for "CODEC enables ‘single duplex’ sequencing"

**Supplementary Table S1. Evaluation of SNP + small indel calls between CODEC WGS and standard WGS. Table are generated by Vcfeval.**

| Method | True-pos-baseline | True-pos-call | False-pos | False-neg | Precision | Sensitivity | F-measure | type | ds_frac | FPPM | FNR | depth |
|---|---|---|---|---|---|---|---|---|---|---|---|---|
| CODEC | 1149572 | 1149620 | 11542 | 2541289 | 0.9901 | 0.3115 | 0.4739 | na12878_cds | 0.1 | 4.482217885 | 0.688526593 | 1 |
| | 1786822 | 1786884 | 17480 | 1904039 | 0.9903 | 0.4841 | 0.6503 | na12878_cds | 0.2 | 6.788179573 | 0.515870691 | 2 |
| | 2222868 | 2222945 | 21154 | 1467993 | 0.9906 | 0.6023 | 0.7491 | na12878_cds | 0.3 | 8.21493997 | 0.397728978 | 3 |
| | 2544500 | 2544593 | 23624 | 1146361 | 0.9908 | 0.6894 | 0.8131 | na12878_cds | 0.4 | 9.174139258 | 0.31058664 | 4 |
| | 2782660 | 2782755 | 25191 | 908201 | 0.991 | 0.7539 | 0.8564 | na12878_cds | 0.5 | 9.782667713 | 0.246061183 | 5 |
| | 2959035 | 2959132 | 25577 | 731826 | 0.9914 | 0.8017 | 0.8865 | na12878_cds | 0.6 | 9.932566873 | 0.198275353 | 6 |
| | 3089997 | 3090099 | 25548 | 600864 | 0.9918 | 0.8372 | 0.908 | na12878_cds | 0.7 | 9.921305019 | 0.162793287 | 7 |
| | 3186894 | 3186995 | 25231 | 503967 | 0.9921 | 0.8635 | 0.9233 | na12878_cds | 0.8 | 9.798201304 | 0.136540826 | 8 |
| | 3259747 | 3259845 | 24540 | 431114 | 0.9925 | 0.8832 | 0.9347 | na12878_cds | 0.9 | 9.529858508 | 0.116802706 | 9 |
| | 3314661 | 3314761 | 23829 | 376200 | 0.9929 | 0.8981 | 0.9431 | na12878_cds | 1 | 9.253748915 | 0.101924675 | 10 |
| Standard NGS | 1362228 | 1362280 | 263618 | 2328633 | 0.8379 | 0.3691 | 0.5124 | na12878_r1r2 | 0.077 | 102.3733594 | 0.630909751 | 1 |
| | 2001169 | 2001242 | 332255 | 1689692 | 0.8576 | 0.5422 | 0.6644 | na12878_r1r2 | 0.15 | 129.0278378 | 0.457795236 | 2 |
| | 2453841 | 2453932 | 366744 | 1237020 | 0.87 | 0.6648 | 0.7537 | na12878_r1r2 | 0.23 | 142.4212889 | 0.335149306 | 3 |
| | 2778411 | 2778517 | 387976 | 912450 | 0.8775 | 0.7528 | 0.8104 | na12878_r1r2 | 0.31 | 150.6665193 | 0.247211639 | 4 |
| | 2990581 | 2990709 | 392938 | 700280 | 0.8839 | 0.8103 | 0.8455 | na12878_r1r2 | 0.38 | 152.5934614 | 0.189726927 | 5 |
| | 3169483 | 3169605 | 380937 | 521378 | 0.8927 | 0.8587 | 0.8754 | na12878_r1r2 | 0.46 | 147.9329955 | 0.141257221 | 6 |
| | 3296709 | 3296823 | 354625 | 394152 | 0.9029 | 0.8932 | 0.898 | na12878_r1r2 | 0.54 | 137.7149989 | 0.106788044 | 7 |
| | 3386823 | 3386934 | 320037 | 304038 | 0.9137 | 0.9176 | 0.9156 | na12878_r1r2 | 0.62 | 124.2831022 | 0.082373424 | 8 |
| | 3443475 | 3443589 | 289759 | 247386 | 0.9224 | 0.933 | 0.9276 | na12878_r1r2 | 0.69 | 112.5249499 | 0.067024567 | 9 |
| | 3491018 | 3491139 | 258701 | 199843 | 0.931 | 0.9459 | 0.9384 | na12878_r1r2 | 0.77 | 100.4638927 | 0.054143586 | 10 |

ds_frac: downsample fraction. FFPM: False positive per million bases. FNR: False negative ratio

**Supplementary Table S2. Sequences of oligonucleotides. Colors match Figure 1.**

| | CODEC Adapter |
|---|---|
| LD4-adap5-1 | AATGATACGGCGACCACCGAGATCTACACCTTGAACGGACTGTCCAC*T |
| LD4-adap5-2 | AATGATACGGCGACCACCGAGATCTACACGAGCCTACTCAGTCAACG*T |
| LD4-adap5-3 | AATGATACGGCGACCACCGAGATCTACACGCTTGTAAGGCAGGTTAG*T |
| LD4-adap5-4 | AATGATACGGCGACCACCGAGATCTACACCAAGCGTCTTACATGGTC*T |
| LD4-adap7-1 | CAAGCAGAAGACGGCATACGAGATCACCGAGCGTTAGACTAC*T |
| LD4-adap7-2 | CAAGCAGAAGACGGCATACGAGATGTGTCGAACACTTGACGG*T |
| LD4-adap7-3 | CAAGCAGAAGACGGCATACGAGATCTGATCTTCAGCTGACTG*T |
| LD4-adap7-4 | CAAGCAGAAGACGGCATACGAGATGAATCTGAGGCACTGTAC*T |
| LD4-brid5-1 | P-GTGGACAGTCCGTTCAAGNNNAGATCGGAAGAGCGTCGTGTAGGGAAAGAGTGTTTACATAGTTATCCGCTAGACTCTGACGTGTTGATCCTCGAAGC |
| LD4-brid5-2 | P-CGTTGACTGAGTAGGCTCNNNAGATCGGAAGAGCGTCGTGTAGGGAAAGAGTGTTTACATAGTTATCCGCTAGACTCTGACGTGTTGATCCTCGAAGC |
| LD4-brid5-3 | P-CTAACCTGCCTTACAAGCTNNNAGATCGGAAGAGCGTCGTGTAGGGAAAGAGTGTTTACATAGTTATCCGCTAGACTCTGACGTGTTGATCCTCGAAGC |
| LD4-brid5-4 | P-GACCATGTAAGACGCTTGANNNAGATCGGAAGAGCGTCGTGTAGGGAAAGAGTGTTTACATAGTTATCCGCTAGACTCTGACGTGTTGATCCTCGAAGC |
| LD4-brid7-1 | P-GTAGTCTAACGCTCGGTGNNNAGATCGGAAGAGCACACGTCTGAACTCCAGTCACCAATCTATAAGTTGCTTCGAGGATCAACACGTCAGAGTCTAGC |
| LD4-brid7-2 | P-CCGTCAAGTGTTCGACACNNNAGATCGGAAGAGCACACGTCTGAACTCCAGTCACCAATCTATAAGTTGCTTCGAGGATCAACACGTCAGAGTCTAGC |
| LD4-brid7-3 | P-CAGTCAGCTGAAGATCAGTNNNAGATCGGAAGAGCACACGTCTGAACTCCAGTCACCAATCTATAAGTTGCTTCGAGGATCAACACGTCAGAGTCTAGC |
| LD4-brid7-4 | P-GTACAGTGCCTCAGATTCANNNAGATCGGAAGAGCACACGTCTGAACTCCAGTCACCAATCTATAAGTTGCTTCGAGGATCAACACGTCAGAGTCTAGC |
| | **CODEC Blocker for Hybridization Capture** |
| LD4-HybBlk1 | AGATCGGAAGAGCGTCGTGTAGGGAAAGAGTGTTTACATAGTTATCCGCTAGACTCTGACGT-3C |
| LD4-HybBlk2 | AGATCGGAAGAGCACACGTCTGAACTCCAGTCACCAATCTATAAGTTGCTTCGAGGATCAAC-3C |

"*" between nucleotides indicates phosphorothioate backbone modification.
"P-" indicates 5'-phosphorylation.
"-3C" indicates C3 spacer.

**Supplementary Table S3. Somatic SNVs of patient 315 found in IDT pan-cancer panel (800kb).**

| Hugo_Symbol | Chromosome | Start_Position | Reference_Allele | Tumor_Seq_Allele2 | tumor_fraction | tumor_ref_count | tumor_alt_count | normal_ref_count | normal_alt_count |
|---|---|---|---|---|---|---|---|---|---|
| PIK3CA | 3 | 178936091 | G | C | 0.245614 | 43 | 14 | 178 | 0 |
| CDKN2A | 9 | 21971000 | C | A | 0.264151 | 39 | 14 | 158 | 1 |
| ARHGAP35 | 19 | 47507737 | C | T | 0.28125 | 46 | 18 | 140 | 0 |
